# Immune checkpoint inhibitor treatment does not impair ovarian or endocrine function in a mouse model of triple negative breast cancer

**DOI:** 10.1101/2024.08.14.607933

**Authors:** Payton De La Cruz, Morgan F. Woodman-Sousa, Julia N. McAdams, Ellia Sweeney, Lola Hakim, Melanie Morales Aquino, Kathryn J. Grive

## Abstract

**Background:** Representing 15-20% of all breast cancer cases, triple negative breast cancer (TNBC) is diagnosed more frequently in reproductive-age women and exhibits higher rates of disease metastasis and recurrence when compared with other subtypes. Few targeted treatments exist for TNBC, and many patients experience infertility and endocrine disruption as a result of frontline chemotherapy treatment. While they are a promising option for less toxic therapeutic approaches, little is known about the effects of immune checkpoint inhibitors on reproductive and endocrine function.

**Results:** Our findings in a syngeneic TNBC mouse model revealed that therapeutically relevant immunotherapies targeting PD-1, LAG-3, and TIM-3 had no effect on the quality and abundance of ovarian follicles, estrus cyclicity, or hormonal homeostasis. Similarly, in a tumor-free mouse model, we found that ovarian architecture, follicle abundance, estrus cyclicity, and ovulatory efficiency remain unchanged by PD-1 blockade.

**Conclusions:** Taken together, our results suggest that immunotherapy may be a promising component of fertility-sparing therapeutic regimens for patients that wish to retain ovarian and endocrine function after cancer treatment.

## INTRODUCTION

Representing 15-20% of all breast cancer cases, triple negative breast cancer (TNBC) is an aggressive form of breast cancer that is diagnosed more frequently in reproductive-age women than other sub-types^1,2^. Because TNBC lacks expression of the estrogen receptor (ER), progesterone receptor (PR), and human epidermal growth factor receptor 2 (HER2), it is unresponsive to many of the targeted treatments that are used for other subtypes of breast cancer^3,4^. Thus, cytotoxic chemotherapies, which are associated with several severe systemic side effects, are often key components of frontline treatments^5^. Indeed, these side effects can impact the reproductive system, and many as half of women who receive conventional chemotherapies experience reproductive and endocrine dysfunction as a result^6^. However, recent advances in cancer immunotherapy have led to the rapid integration of Pembrolizumab, a programmed cell death protein 1 (PD-1) inhibitor, into standard-of-care treatment regimens for TNBC, which may allow for reduced toxicity during treatment^7,8^.

The development of immunotherapies has shown great promise for the targeted treatment of a variety of cancers, and they lack many of the side effects observed with standard of care chemotherapeutics^9^. One such class of immunotherapies that have seen clinical success is immune checkpoint inhibitors, which target immune checkpoint regulators such as PD-1 and its ligand PD-L1^5^. These ligands act as modulators in normal tissue to promote self-tolerance but are overexpressed in tumor cells as a mode of evading immunosuppression^10^. Immune checkpoint inhibitors such Pembrolizumab employ monoclonal antibodies to block immune checkpoint interactions with the goal of activating tumor-specific cytotoxic T cells and promoting immune cell-mediated killing^4^.

Though the introduction of immune checkpoint inhibitors has revolutionized cancer treatment paradigms and patient quality of life, the systemic blocking of immune checkpoint interactions can cause immune-related adverse events (irAEs) associated with a lack of immune tolerance of self-tissues^11^. While these effects are not as common as those associated with cytotoxic chemotherapy and are usually mild, they can occasionally be severe and may affect a variety of organs systems^9^. Indeed, endocrine irAEs are some of the most commonly reported irAEs in the clinic and include hyper- and hypothyroidism, hypophysitis, hypogonadism, and Type I diabetes^12,13^. In a 2022 study by Winship et al., anti-CTLA-4 and anti-PD-L1 monoclonal antibodies were associated with an increase in intra-ovarian immune cells and tumor necrosis factor-α (TNF-α) expression, as well as a decrease in ovarian follicle quality and abundance^14^. However, no studies have evaluated the ovarian and endocrine effects of standard-of-care PD-1 inhibitors or blockade of other exploratory targets such as lymphocyte-activation gene 3 (LAG-3) or T cell immunoglobulin and mucin domain 3 (TIM-3).

In the mammalian ovary, immature oocytes are stored in a quiescent state as primordial follicles in a finite population known as the ovarian reserve^15,16^. After puberty, these primordial follicles are continuously activated to begin folliculogenesis, the process of transforming into larger, mature follicles that produce steroid hormones and may eventually be ovulated^17,18^. This process occurs continuously through the entire reproductive lifespan until the ovarian reserve is depleted, at which point menopause begins^19,20^. Because new oocytes cannot be generated after birth, it is critical that primordial follicles must be available in sufficient amounts and be maintained through the reproductive lifespan of the host to ensure fertility and endocrine function^21,22^. Cytotoxic chemotherapies are among the foremost causes of ovarian reserve damage, often resulting in the condition of Primary Ovarian Insufficiency (POI)^6,23^. Caused by the depletion of the ovarian reserve, POI can lead to infertility and impaired endocrine function, and increases the risk of conditions associated with aging such as heart attack, stroke, and osteoporosis^24^.

In all mammals, ovarian follicles undergo highly-conserved processes of growth, maturation, ovulation, and death, although the timeline of these events is species-specific^16,17^. In humans, the time for an activated primordial follicle to mature into a fully-grown pre-ovulatory follicle takes about 12 months, while in mice, this process takes around 21 days^25,26^. The menstrual cycle in humans takes an average of 28 days and typically results in the ovulation of a single oocyte per cycle, while the mouse estrous cycle takes only 4-5 days and results in the ovulation of several oocytes per cycle^19,26^. The mouse reproductive cycle begins around 6-8 weeks of age and ends approximately after 12 months^26,27^. In humans, the menstrual cycle begins at the onset of puberty and continues until menopause^16,19^. Though mice do not undergo a true “menopause”, they experience similarities to human females in the processes of ovarian reserve depletion, loss of fertility, and endocrine dysfunction with aging, and are therefore tremendously useful models for study of mammalian reproductive function^28^.

Immune regulation plays an important role in reproductive and endocrine health. The ovary is subject to immune cell infiltration as it is a highly vascularized, non-immunoprivileged organ^29^. In the ovary, immune cells are critical in processes related to granulosa cell turnover, ovulation, clearing atretic follicles, and the development and breakdown of the corpus luteum^30,31^. In addition, signaling by cytokines such as TNF-α and transforming growth factor-β (TGF-β) are crucial in follicle maturation and ovulation^20^. However, it is likely that dysregulation or over-activation of these immune factors could cause damage to the ovary^32–34^. A few epidemiologic studies have reported a higher incidence of POI and unexplained infertility in women with chronic inflammatory or autoimmune conditions such as Crohn’s Disease and psoriasis^35,36^. In addition, there is some evidence linking inflammation and autoimmunity on ovarian aging and follicle depletion in animal models^34^. However, the specific mechanisms of autoimmune depletion of the ovarian reserve remain unknown, and it is unclear whether immune checkpoint inhibitor treatment creates a sufficiently heightened systemic immune response to elicit ovarian damage and endocrine disruption.

The ovarian toxicity of many of the frontline chemotherapeutics for TNBC has been well-characterized, with the non-renewable population of oocytes being some of the most vulnerable cells to damage^6,23^. As immune checkpoint inhibitors continue to be incorporated into clinical practice, more studies on their reproductive and endocrine effects are necessary, especially if they are to be used as a component of fertility-sparing treatment regimen. Moreover, given the fact that they are still relatively new to the cancer therapeutic repertoire, data on long-term fertility outcomes in human TNBC patients treated with immune checkpoint inhibitors is not yet available and thus preclinical models must be used to evaluate their ovarian impacts^37^. We hypothesized that ICIs targeting PD-1, LAG-3 and TIM-3 would be relatively benign to ovarian reserve and endocrine function.

## MATERIAL AND METHODS

### Animals

Wild-type C57Bl/6 mice were obtained from the Jackson Laboratory (strain # 000664). All animal protocols were approved by Brown University Institutional Animal Care and Use Committee and were performed in accordance with the National Institutes of Health Guide for the Care and Use of Laboratory Animals (# 22-09-0002). All animal protocols were reviewed and acknowledged by the Lifespan University Institutional Care and Use Committee (# 1987412-1).

### E0771 tumor-bearing mouse model and tissue collection

Mouse E0771 cells were obtained from ATCC and cultured in DMEM, 10% FBS, and penicillin/streptomycin. Cells were found to be free of pathogens and mycoplasma (per Charles River pathogen testing). Eight-week-old female C57Bl/6J mice were injected with 100 μL of 1×10^5^ E0771 cell suspension in Matrigel or saline control into the bilateral 4^th^ mammary pads under isoflurane sedation. Once palpable 14 days later, a group of pre-treatment mice were collected, and remaining mice were randomly allocated into study groups to begin treatment regimen of immune checkpoint inhibitor monotherapy or control. Mice received 200 μg doses of mouse anti-PD-1 (clone: 29F.1A12), anti-LAG-3 (clone: C9B7W), anti-TIM-3 (clone: RMT3-23), or rat IgG2a isotype control (clone: 2A3) every 4 days via intraperitoneal injection, with treatments stopping after the third dose. All antibodies used for *in-vivo* treatments were purchased from BioXcell. Doses were based on previously described tumor-reducing regimens^45^. Mice were monitored for 14 days and then collected once they reached proestrus stage. Upon collection, tumors, ovaries, and serum were obtained for further analysis. Tumor burden was determined by quantifying tumor weight as a proportion of total mouse weight.

### Non-tumor-bearing mouse model and tissue collection

Eight-week-old female C57Bl/6J mice were mock-injected with 100 μL saline in the 4^th^ mammary pad under isoflurane sedation. 14 days later, mice were randomly allocated into study groups to begin treatment with anti-PD-1 monoclonal antibody, IgG isotype control, or saline via intraperitoneal injection. Mice in the anti-PD-1 and IgG isotype control group received 200 μg doses of monoclonal antibodies every 4 days via intraperitoneal injection, with treatments stopping after the third dose. Mice were monitored for 14 days and then collected once they reached proestrus stage. Upon collection, ovaries and serum were obtained for further analysis.

### Vaginal cytology and estrus cycle analysis

After the final dose of immunotherapy or control, estrus cycle stage was monitored daily over the course of 14 days via vaginal smear cytology as previously described^46^. Briefly, the vagina of each mouse was flushed with saline and then mixed with toluidine blue O dye on a glass slide, then classified into the different sub-stages of estrus based on the cell types visualized in the sample. The percentage of time spent in each sub-stage of estrus was calculated for each mouse and results were compared between treatment groups.

### Ovarian histology and follicle quantification

Ovaries from tumor-bearing and non-tumor-bearing mice were stained and analyzed as previously described^46^. Briefly, ovaries were fixed in formalin and embedded in paraffin for sectioning at 5 μm by the Brown Molecular Pathology Core, and every fourth slide was deparaffinized and stained with hematoxylin and eosin (H&E). Five slides per ovary were quantified. Follicles were staged and counted on one section per every H&E-stained slide, and these counts were normalized to section area to yield follicle density.

### TUNEL staining

Prior to staining, ovarian FFPE slides were deparaffinized as previously described. Slides were then washed for 3 minutes in Phosphate Buffered Saline (PBS), then permeabilized by applying freshly prepared 20 μg/mL Proteinase K in 10mM Tris-HCl solution and incubating for 15 minutes in a humid chamber at room temperature. Slides were then washed 2x, 3 minutes in PBS, 1x 3 minutes in PBS + 0.1% Triton (PBST), then again 2x, 3 minutes in PBS. Slides were then stained with the *In Situ Cell Death Detection Kit, Fluorescein* (Roche) according to manufacturer’s instructions. Briefly, slides were incubated with TUNEL reaction mixture for 1 hour in a humid chamber at 37°C protected from light. Slides were then washed 3x for 3 minutes in PBS, once for 3 minutes in DAPI/PBS solution, and then once for 5 minutes in PBS. Slides were then mounted and analyzed on an EVOS M5000 Fluorescence Imaging System and images of all fields of a single section were captured.

### Serum hormone analysis

Whole blood from mice was collected via post-mortem cardiac puncture and serum separated out as previously described^46^. Serum was sent to the University of Virginia Ligand Assay & Analysis Core for the Center for Research in Reproduction for quantification of serum concentrations of AMH, LH, and FSH.

### Superovulation of tumor-bearing and non-tumor-bearing mice

For superovulation experiments in tumor-bearing mice, eight-week-old wild-type C57Bl/6J mice were orthotopically injected with E0771 cells into the right 4^th^ mammary pad as previously described. Once palpable 14 days later, mice were randomly allocated into groups to receive intraperitoneal injections of anti-200 μg doses of PD-1 monoclonal antibody or IgG isotype control, or 100 μL of saline control every 4 days with treatments stopping after the third dose. Eight days after receiving their final dose, mice underwent ovarian hyperstimulation as previously described^46^. Briefly, mice were injected with 5-IU pregnant mare goat serum (PMSG; Prospec Bio), and then 48 hours later, injected with 5-IU human chorionic gonadotropin (HCG; Prospec Bio). Twelve hours after HCG injection, mice were euthanized, and ovulated oocytes were collected from the ampullae. Number of oocytes ovulated were counted per mouse and results were compared between treatment groups.

For superovulation experiments in non-tumor-bearing mice, six-week-old wild-type C57Bl/6J mice were mock-injected with saline into the mammary pad as described. As with the tumor-bearing superovulation study, mice were randomly allocated into groups 14 days later to receive intraperitoneal injections of anti-200 μg doses of PD-1 monoclonal antibody or IgG isotype control, or 100 μL of saline control every 4 days with treatments stopping after the third dose. Eight days after receiving their final dose, mice underwent ovarian hyperstimulation and ovulated oocytes were quantified as described.

### Statistical analysis

Ovarian area and follicle density were quantified from H&E-stained ovarian sections. Follicles were counted by stage, and these counts were then normalized to the section area to calculate oocyte density and account for size differences between different ovaries. For estrus cycling analyses, the percentage of time spent in each substage was then averaged between all animals in a treatment group. For superovulation studies, recovered oocytes were quantified and averaged per treatment group, and mice were classified as “successfully stimulated” if the number of retrieved oocytes met an age-adjusted threshold based on average litter sizes for wild-type C57Bl/6J mice in our lab (7 oocytes for 8-week-old mice, 10 oocytes for 6-week-old mice). One-way ANOVA with post-hoc Tukey’s tests for multiple comparisons were performed to evaluate differences in tumor burden, follicle abundance, serum hormone levels, percentage of time spent in estrus between treatment groups.

## RESULTS

### Inhibition of PD-1, LAG-3, and TIM-3 does not impact ovarian follicle abundance or quality in a mouse model of triple negative breast cancer

To assess the effects of immune checkpoint inhibition on ovarian health and endocrine function in a clinically-relevant model of TNBC, we collected ovaries and serum of C57Bl/6J mice who had been orthotopically injected with syngeneic E0771 cells into the mammary pad, then treated with intraperitoneal injections of anti-PD-1, anti-LAG-3, or anti-TIM-3 monotherapy. These immunotherapies represent an array of immune checkpoint inhibitor candidates that have demonstrated varying efficacy in clinical trials for solid cancers, with anti-PD-1 being the most effective in TNBC populations^38–40^. By choosing these specific targets, we were able to comprehensively evaluate the effects of differential immune activation on the ovarian reserve and hormonal homeostasis. Tumor-bearing control group mice received intraperitoneal injections of IgG isotype at the same timepoints as immunotherapy-treated mice, and healthy control mice only received a mock-injection of saline into the mammary pad at the time of orthotopic injections. To control for cycle stage-dependent fluctuations in folliculogenesis, ovaries were collected during the proestrus stage 14-16 days following the final immunotherapy treatment. This collection timepoint allowed us to capture any effects that may have a pattern of delayed onset seen in some other adverse immune effects.

Morphologically, ovarian architecture was similar among all treatment groups (Fig 1a-f). Follicles were classified by stage, including the immature, quiescent primordial follicles that make up the ovarian reserve, the developing primary, secondary, and preantral follicles that have been recruited and activated to undergo maturation, and the antral follicles that are nearly ready for ovulation. Degenerative follicles, which are undergoing atresia and dying, were also quantified per section. At the post-treatment timepoint, anti-PD-1-treated mice showed almost complete tumor regression, which was statistically significant when compared to the IgG isotype control group (p=0.0125) (Fig 1f). Mice treated with anti-LAG-3 or anti-TIM-3 exhibited a more variable response to treatment, with the anti-TIM-3-treated group achieving a significant reduction in tumor burden (p=0.0492). Though the anti-LAG-3 group had a lower mean tumor burden than IgG isotype controls, this difference was not statistically significant (p=0.8006). Ovarian area, as well as overall and stage-specific oocyte densities were not significantly different between immune checkpoint inhibitor-treated groups and IgG isotype-treated or saline mock-injected controls (Fig 1g-i). There were also no appreciable levels of oocyte or granulosa cell apoptosis found via TUNEL staining (Supp Fig 1). These findings indicate that inhibition of a variety of immune checkpoint interactions has no long-term effect on ovarian morphology or folliculogenesis in a tumor-bearing system.

**Figure 1.**
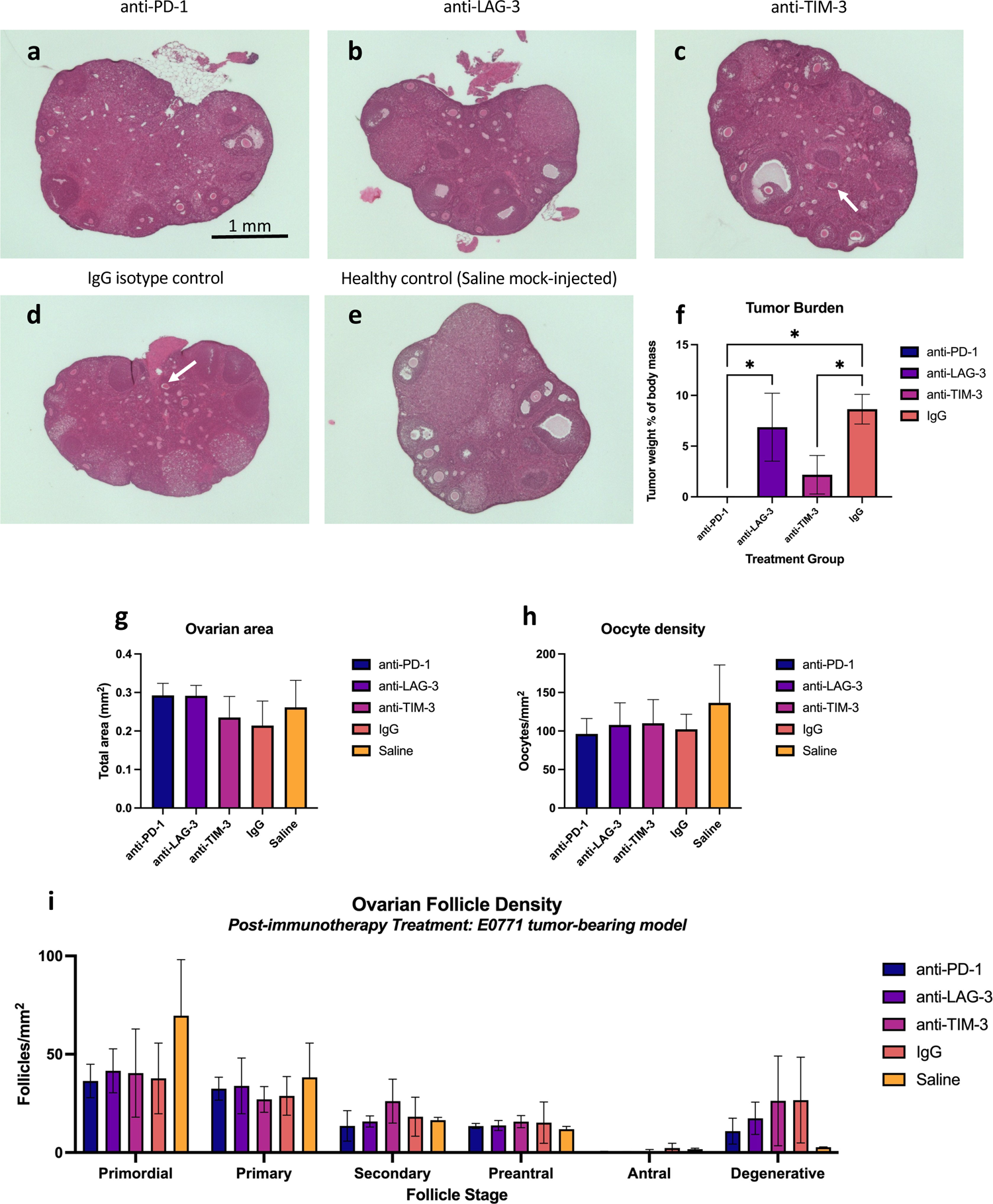
Inhibition of PD-1, LAG-3, and TIM-3 does not impact ovarian follicle abundance or quality in a mouse model of triple negative breast cancer. Ovaries from E0771 tumor-bearing mice treated with monoclonal antibodies targeting PD-1, LAG-3, TIM-3 (**A-C**), IgG isotype (**D**), and a healthy control mock-injected with saline (**E**) were collected, formalin-fixed, paraffin-embedded, and then stained with hematoxylin and eosin via standard protocols (n=3 per group). Degenerative follicles are denoted with white arrows. Tumor burden analysis (**F**) at study endpoint reveals near-complete tumor regression in anti-PD-1-treated mice, along with varying reductions in tumor size in the anti-LAG-3 and anti-TIM-3 groups (n=3 per treatment group). Ovarian size, oocyte density, and follicle stage density was not significantly different between any of the treatment or control groups (**G-I**). Ovarian follicle counts were quantified using one section on every fourth slide of sectioned ovary to capture all follicle stages without over-representing larger follicles. To account for size differences between ovaries, section area was used to normalize follicle counts. Data for follicle counts, follicle density, and ovarian area can be found in Supplementary File 1. *p<0.05, as indicated,

### Inhibition of PD-1, LAG-3, or TIM-3 does not perturb endocrine homeostasis or reproductive cyclicity

Given that long-term endocrine dysfunction and disruption of reproductive cycling are common side effects of cancer therapy that are at times unrelated to ovarian reserve function, we assessed these outcomes via serum hormone and estrus cycling analysis in the E0771 tumor-bearing mouse model. As with the ovarian collections, we collected serum during the proestrus stage 14-16 days following the final immunotherapy treatment to control for cycle stage-dependent fluctuations in hormone levels. Serum levels of luteinizing hormone (LH) and follicle stimulating hormone (FSH) were quantified to evaluate possible disturbances in the hypothalamic-pituitary-gonadal axis and hormonal cycling. Serum levels of anti-Mullerian hormone (AMH), produced by granulosa cells of maturing follicles and current clinical gold standard for evaluating follicle abundance, was also quantified^41^. Serum levels of LH and FSH levels did not differ significantly between any of the immunotherapy treatment and control groups (Fig 2a-b). Likewise, there were no significant differences observed in LH:FSH ratio (Fig 2c), a common clinical metric in which high values correlate with a lack of ovulation in polycystic ovarian syndrome^42^. Consistent with our ovarian follicle density results, we found that AMH levels were unchanged between immunotherapy-treated and control group animals.

**Figure 2.**
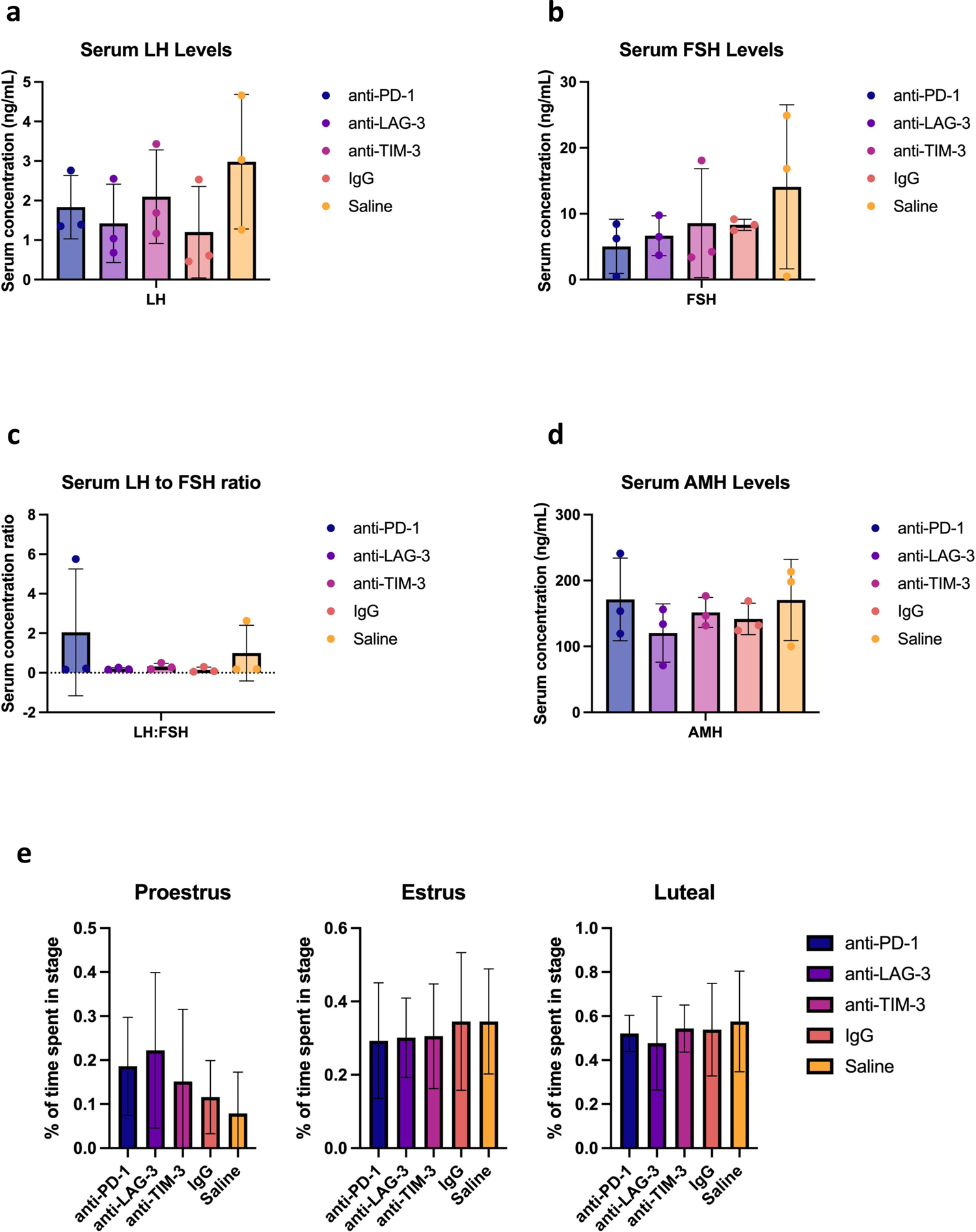
Inhibition of PD-1, LAG-3, and TIM-3 does not perturb endocrine homeostasis or reproductive cyclicity. Serum concentrations of LH (**A**) and FSH (**B**), as well as the LH:FSH ratio (**C**), and AMH (**D**) were not significantly different between any of the treatment and control groups (n=3 per group). Estrus cyclicity was monitored daily via vaginal cytology for two weeks starting the day following the final immunotherapy treatment, and the percentage of time spent in each substage was averaged between mice in each treatment group (n=3 per group). The amount of time spent in each substage of the estrus cycle was unchanged between all treatment and control groups (**E**).

Estrus cyclicity was analyzed by monitoring vaginal cytology daily for two weeks starting the day after the final immunotherapy or IgG isotype control treatment. We found that there were no significant differences in the amount of time spent in each of the cycle stages between any of the immunotherapy treatment and control groups (Figure 2e). Taken together with the serum hormone data, these findings suggest that immune checkpoint blockade does not impair endocrine function or hormonal cyclicity in tumor-bearing mice.

### Treatment with anti-PD-1 immunotherapy does not impact ovarian follicle density or reproductive cyclicity in a non-tumor-bearing mouse model

To further investigate ovarian and endocrine function after treatment with anti-PD-1 immunotherapy and to disentangle any observed effects from tumor burden-specific phenomena, we sought to validate our results in a tumor-free mouse model. To simulate orthotopic implantation of tumor cells, young adult female C57Bl/6J mice age-matched to the E0771 cohort were mock-injected with saline into the mammary pad. Narrowing down our treatment focus to answer questions of key clinical relevancy, we then administered anti-PD-1 mAb, IgG isotype control, or saline control intraperitoneal injections, all at the same timepoints as the cohort of E0771 tumor-bearing mice. As with all tumor-bearing studies, ovaries and serum collected 14-16 days following the final treatment when mice entered the proestrus stage.

Ovaries from all groups of non-tumor-bearing mice appeared morphologically similar to each other (Fig 3a-c) and showed no differences in total area (Fig 3d). Upon quantifying ovarian follicle abundance, anti-PD-1-treated mice did not differ significantly in oocyte and follicle stage densities compared to IgG isotype and saline-treated controls (Fig 3e-f). These findings recapitulate our results from the tumor-bearing model and suggest that PD-1 blockade has no negative effect on follicle abundance or folliculogenesis.

**Figure 3.**
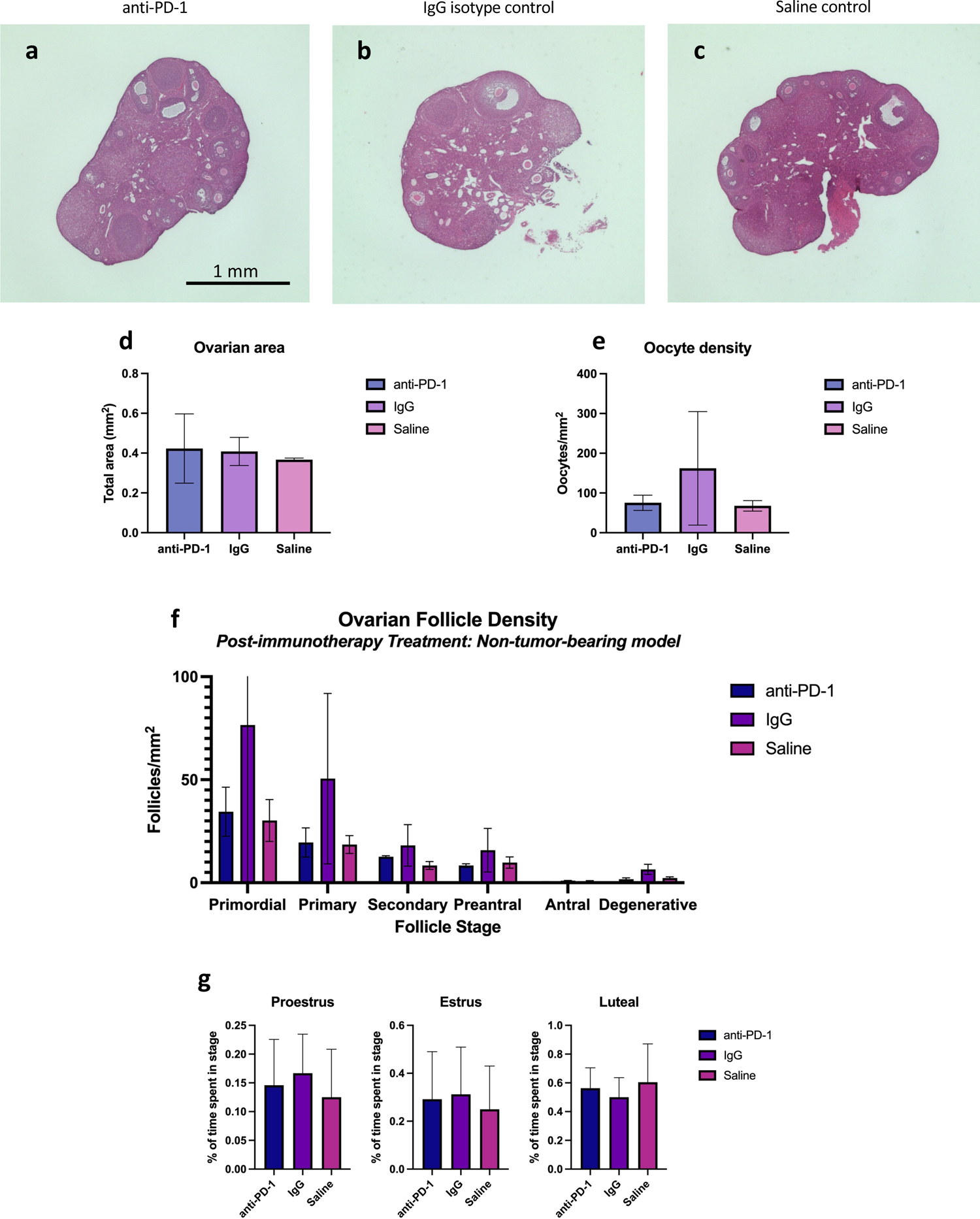
Treatment with anti-PD-1 immunotherapy does not impact ovarian follicle density or reproductive cyclicity in a non-tumor-bearing mouse model. Ovaries were collected for FFPE from healthy, non-tumor-bearing mice treated with anti-PD-1 monoclonal antibodies, IgG isotype control, or saline control (**A-C**), and then stained with hematoxylin and eosin (n= 3 per group). There were no significant differences between ovarian size, oocyte density, follicle abundance, or the percentage of time spent in each substage of estrus between treatment groups (**D-G**). Data for follicle counts, follicle density, and ovarian area can be found in Supplementary File 2.

To evaluate any effects of anti-PD-1 immunotherapy on hormonal cyclicity in the absence of tumor burden, we monitored estrus cycling via vaginal cytology for two weeks following administration of the final immunotherapy or control treatment. Unsurprisingly, we found that the percentage of time spent in each substage of estrus was unchanged between anti-PD-1-treated animals and control animals treated with IgG isotype or saline (Fig 3g). These results are consistent with the tumor-bearing model, suggesting that hormonal cyclicity is unaffected by PD-1 inhibition in both a tumor-bearing and non-tumor-bearing system.

Interestingly, the only differences observed in ovarian composition are found when comparing treatment-matched groups from the tumor-bearing and non-tumor-bearing cohorts. IgG isotype-treated animals from the tumor-bearing group have a statistically significant decrease in ovarian area compared to their non-tumor-bearing counterparts (Supp Fig 2). Likewise, tumor-bearing anti-PD-1-treated animals had higher levels of preantral follicle density than the non-tumor-bearing anti-PD-1-treated group (Supp Fig 2). These findings demonstrate that the effect of tumor burden may play a larger role in determining ovarian and endocrine health than any immune-related changes resulting from PD-1 blockade.

### Treatment with anti-PD-1 immunotherapy does not impair ovulatory capacity

To investigate whether PD-1 inhibition could affect ovulatory efficiency, we conducted superovulation studies to assess responses to ovarian hyperstimulation. One week following treatment with anti-PD-1 immunotherapy, IgG isotype, or saline control, animals of each group were hormonally stimulated with PMSG, then given a trigger shot of HCG 48 hours later. Cumulus-oocyte complexes and ovaries were collected from the ampullas of all animals 12 hours after the HCG injection. Ovaries from stimulated mice displayed corpora lutea formation consistent with recent ovulation (Fig 4 a-c).

**Figure 4.**
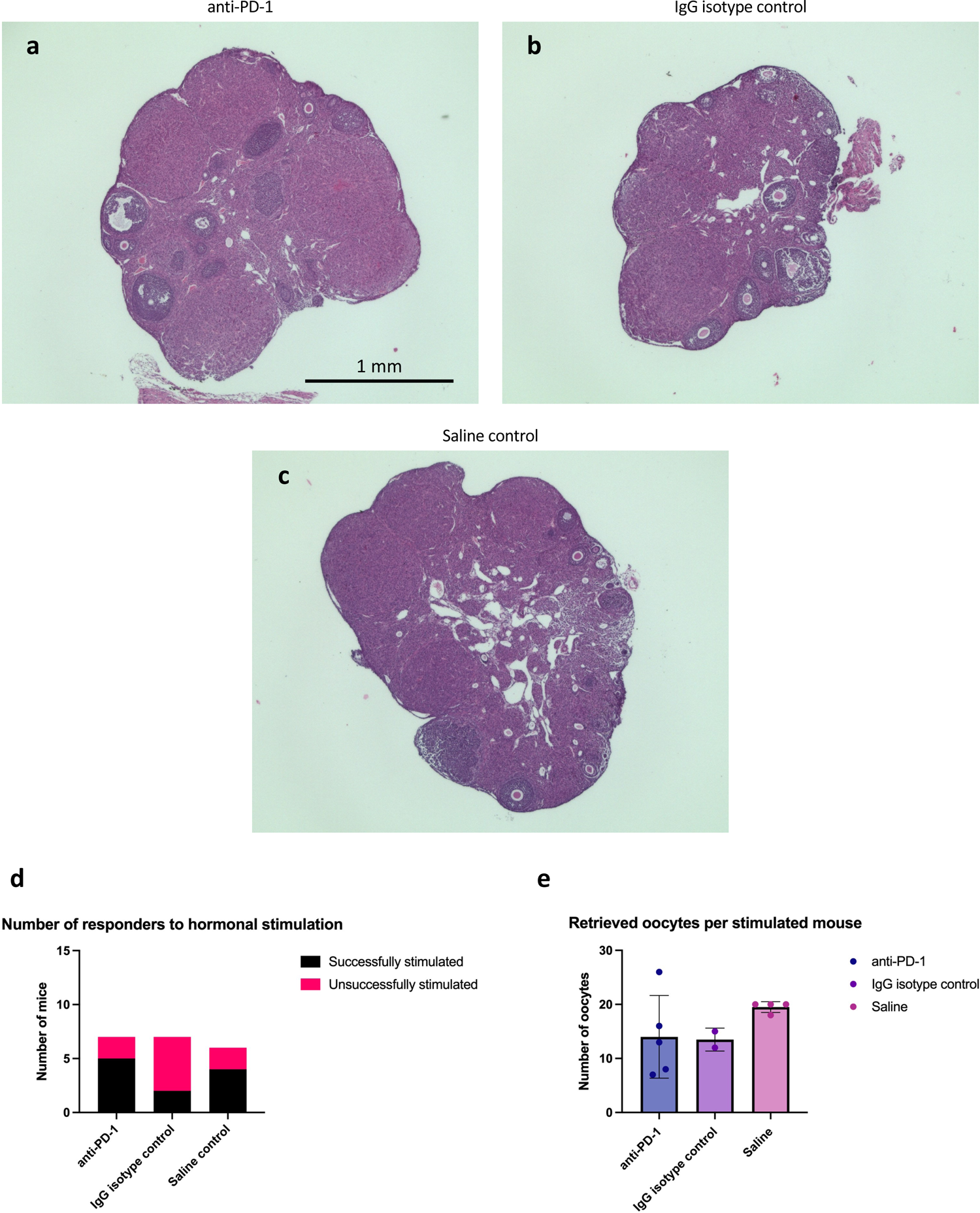
Treatment with anti-PD-1 immunotherapy does not impair ovulatory capacity. One week following treatment with anti-PD-1 monoclonal antibodies, IgG isotype control, or saline control, mice were super-ovulated with PMSG and HCG using standard protocols. Ovaries from superovulated mice were collected for FFPE and then stained with hematoxylin and eosin (**A-C**). There were no significant differences in the number of mice who responded to hormonal stimulation (**D**) or the number of oocytes retrieved from successfully superovulated mice (**E**). Retrieved oocytes were collected from each animal, then counted and averaged per treatment group (n=6 for anti-PD-1 and IgG isotype control, n=5 for saline control). Mice were classified as “successfully stimulated” if their number of retrieved oocytes reached an age-adjusted threshold for hyperstimulation (10 oocytes for 7-week-old mice, 7 oocytes for 10-week-old mice). Counts of retrieved oocytes from “successfully stimulated” mice were averaged and compared between treatment groups (n=5 for anti-PD-1, n=2 for IgG isotype control, n=4 for saline control).

There were no significant differences in the numbers of successfully stimulated mice or number of oocytes recovered per stimulated mouse between the anti-PD-1-treated group and control groups (Fig 4d-e). These results indicate that PD-1 inhibition does not impact ovulatory efficiency or the ability to respond to hormonal stimulation.

## DISCUSSION

As cancer therapeutics and survivorship outcomes improve, preserving the ovarian reserve during cancer treatment has become critical. Not only does the ovarian reserve comprise an individual’s entire reproductive potential, but it also directly contributes to endocrine balance, making it critical for female health-span and lifespan. Moreover, though immune checkpoint inhibitors may potentially be a less-toxic approach to treating cancer, they are still associated with a suite of adverse immune effects and thus require characterization of their impact on reproductive and hormonal function^2,13^. Based on our rigorous studies in both a TNBC tumor-bearing model and a tumor-free model, we found no evidence that immune checkpoint inhibition negatively affects the ovarian reserve or endocrine homeostasis.

In this work, we demonstrate that ovarian architecture and follicle density remain unchanged by immune checkpoint blockade. Ovaries from tumor-bearing mice treated with immunotherapies targeting PD-1, LAG-3, and TIM-3 appeared healthy and bore remarkable resemblance to ovaries from mice treated with IgG isotype control. While we were interested in understanding any ovarian or endocrine effects of anti-PD-1 therapy as it is commonly being used to treat TNBC in the clinic, we chose to also investigate anti-LAG-3 and anti-TIM-3 because of their possible future uses in treatment regimens for TNBC or other solid cancers. However, due to their variable efficacy in TNBC clinical trials, we narrowed our scope to focus solely on anti-PD-1 therapy in non-tumor-bearing and subsequent studies^38^. As seen in the tumor-bearing cohort, ovaries from non-tumor-bearing mice treated with anti-PD-1 were equally as healthy as the control group ovaries. Critically, this lack of ovarian and endocrine damage observed after immune checkpoint inhibitor treatment coincides with tumor regression in the tumor-bearing cohort, particularly in anti-PD-1-treated mice. This anti-tumor efficacy indicates that our monotherapy doses are therapeutically relevant, and thus, are appropriate doses for evaluating reproductive toxicity.

Interestingly, the primary differences were found only when comparing ovaries from tumor-bearing mice to those from non-tumor-bearing mice. Among mice treated with anti-PD-1 therapy, preantral follicles were significantly higher in the tumor-bearing group, while non-significant increases could also be seen in degenerative and primary follicles of the tumor-bearing group. One possible reason for this could be a delay in follicle maturation in tumor-bearing animals, causing an accumulation of maturing follicles and thus a higher percentage of degeneration. These disruptions in folliculogenesis may be attributable to non-specific inflammation related to tumor burden, especially considering the anatomical proximity of the mammary tumor to the ovaries. Moreover, among mice treated with IgG isotype, ovarian area was reduced in the tumor-bearing group while oocyte density remained comparable. This finding could potentially be due to cancer-related inflammatory effects on the somatic compartment of the ovary, a mechanism proposed by Chaqour et al^43^. These results may possibly indicate a trend that local cancer-associated inflammation may have a larger role in determining ovarian health than immune checkpoint blockade itself.

Our studies also show that hormonal cyclicity and serum hormone levels are similar between immunotherapy and control group mice. These results were not particularly surprising given that we found that immunotherapy did not diminish the ovarian reserve. However, considering that autoimmune disorders affecting endocrine function are among the most common of the adverse immune effects associated with immune checkpoint inhibitor treatment, it was important for our research to investigate possible extra-ovarian effects^13^. Finally, our results suggest that anti-PD-1 immunotherapy has no effect on responsiveness to hormonal stimulation or ovulatory capacity. Taken together, we can conclude that immune checkpoint inhibitors, namely anti-PD-1-based immunotherapies, could be a critical component of ovary-sparing cancer therapeutic regimens.

Though there is little literature available on the effects of immunotherapy on the ovarian reserve, a 2022 study from Winship et al. found that PD-L1 and CTLA-4 inhibition causes depletion of the ovarian reserve, disruption of estrus cyclicity, reduced ovulatory capacity, and an increase in intra-ovarian immune activity^14^. Similar to our studies, Winship et al. evaluated these immunotherapies in both a tumor-bearing and non-tumor-bearing model. Given that we investigated different immune checkpoint targets, our results are not necessarily inconsistent with those reported by Winship et al. However, our conclusions are strikingly different when generalizing about the effects of immune checkpoint blockade as a whole. While the Winship study posits that immune checkpoint inhibitors are harmful to the ovarian reserve, our study is the first to test the effects of anti-PD-1 therapy, the only standard-of-care immunotherapy approved by the FDA for the treatment of TNBC^7^. In addition to the difference in selection of immune checkpoint targets, some of our contrasting results may also be attributable to the difference in mouse models used. For example, the Winship study employed the use of C57Bl/6J mice orthotopically injected with AT3OVA cells for their tumor-bearing experiments^14^. While this is indeed a syngeneic model for mammary carcinoma, the cell line is not as commonly used as our E0771 model and could have critical differences in immunogenicity, receptor expression, or clinical relevance^44^. Ultimately, our study provides critical context to the current body of knowledge on the reproductive and endocrine impact of immune checkpoint inhibitors.

Our mouse studies sought to model a clinically-relevant dosing schedule equivalent to one round of immunotherapy treatment widely used in previous research. By allowing 14 days to elapse between treatment and collection, we were able to monitor and record daily estrus cycling to assess immediate effects before eventually capturing delayed effects on ovarian health and serum hormone balance. Future studies will target long-term outcomes such as fecundity and premature ovarian aging after immunotherapy exposure.

Though we aimed to design and execute comprehensive studies, this work is not without its limitations. Our studies do not investigate subtle immunotherapy-induced changes that may not be apparent in histological analyses. However, due to the overwhelming lack of negative functional effects observed, any subtle phenotypes will be subject of future studies and are beyond the purview of this current study. In addition, we recognize that the number of mice who were responsive to hormonal stimulation is low, making it more difficult to draw conclusions about ovulatory capacity. However, considering the need to balance a clinically-relevant dosing schema with the inherent challenges of hormonally stimulating mice older than 4 weeks of age, we chose to perform our study as described and make statistical adjustments to account for reasonable oocyte recoveries.

## CONCLUSIONS

Our research adds novel information to a burgeoning field studying the effects of immunotherapy on reproductive and endocrine function, and provides reassuring data to support that anti-PD-1 therapy for TNBC, which is now standard-of-care, does not detrimentally affect ovarian function. Given the rising number of young women being diagnosed with TNBC, there is a critical need to design treatment approaches that can maximize therapeutic efficacy while minimizing damage to the ovarian reserve, thereby improving treatment outcomes and quality of life for survivors. Though population-level clinical data would provide the most clarity on the fertility outcomes after immune checkpoint inhibitor treatment, our studies present a human-relevant animal model to assess ovarian and endocrine effects of novel immunotherapies to inform clinical recommendations.

## DECLARATIONS

### Ethics approval and consent to participate

Not applicable.

### Consent for publication

Not applicable.

### Availability of data and materials

The data generated from this study are available from the corresponding author upon reasonable request.

### Authors’ contributions

PDLC, MFW-S, JNM, ES, LH, MMA, and KJG performed the experiments. PDLC and KJG analyzed and interpreted the data. PDLC and KJG wrote and revised the manuscript.

## Acknowledgements

The authors would like to thank the Program in Women’s Oncology of Women and Infants Hospital and the generous funding support of Swim Across America and the Foundation for Women’s Wellness. Technical support for this work was provided by the University of Virginia Center for Research in Reproduction Ligand Assay and Analysis Core, which is supported by the Eunice Kennedy Shriver NICHD Grant R24 HD102061. The authors thank the Freiman and James laboratories for providing valuable feedback on this project.

---

***Supplemental* figure 1** TUNEL staining of ovaries from E0771 tumor-bearing mice treated with monoclonal antibodies targeting PD-1, LAG-3, TIM-3, and IgG isotype control showed no appreciable levels of apoptosis in follicles.

***Supplemental* figure 2** Differences in ovarian area, follicle density, and estrus cyclicity in tumor-bearing vs. non-tumor-bearing mice. Among mice treated with IgG isotype control, ovarian area was significantly increased in non-tumor-bearing mice, which may indicate increased volume in the ovarian somatic compartment of healthy mice. Among mice receiving anti-PD-1 treatment, preantral follicle counts were significantly higher in the tumor-bearing group, perhaps indicating an accumulation of maturing follicles in TNBC mice. *p<0.05, as indicated

