## Supplementary figures and images for "Immune checkpoint inhibitor treatment does not impair ovarian or endocrine function in a mouse model of triple negative breast cancer"

### Supp Figs 1 and 2

*Supplemental Figure 1*

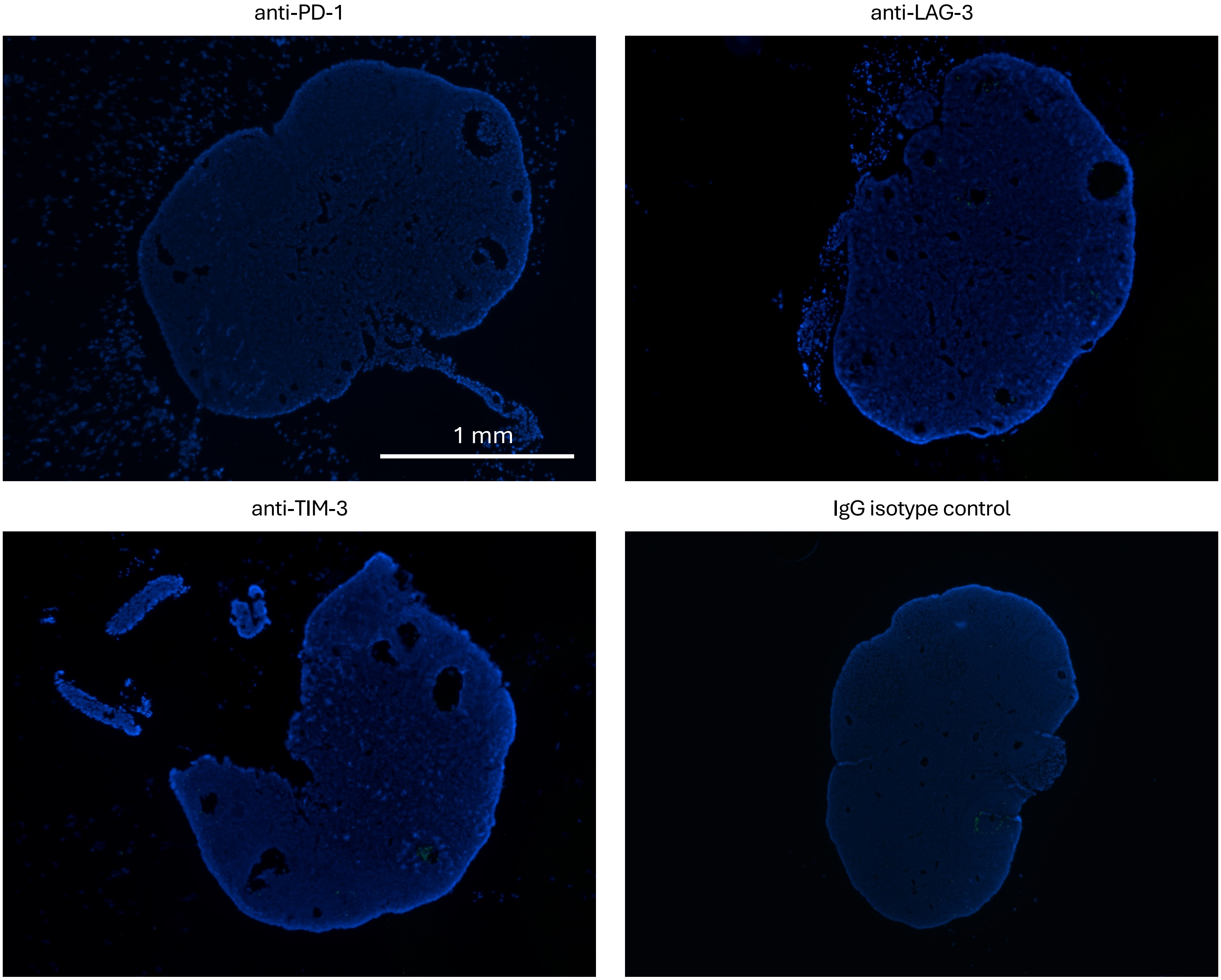

*Supplemental Figure 2*

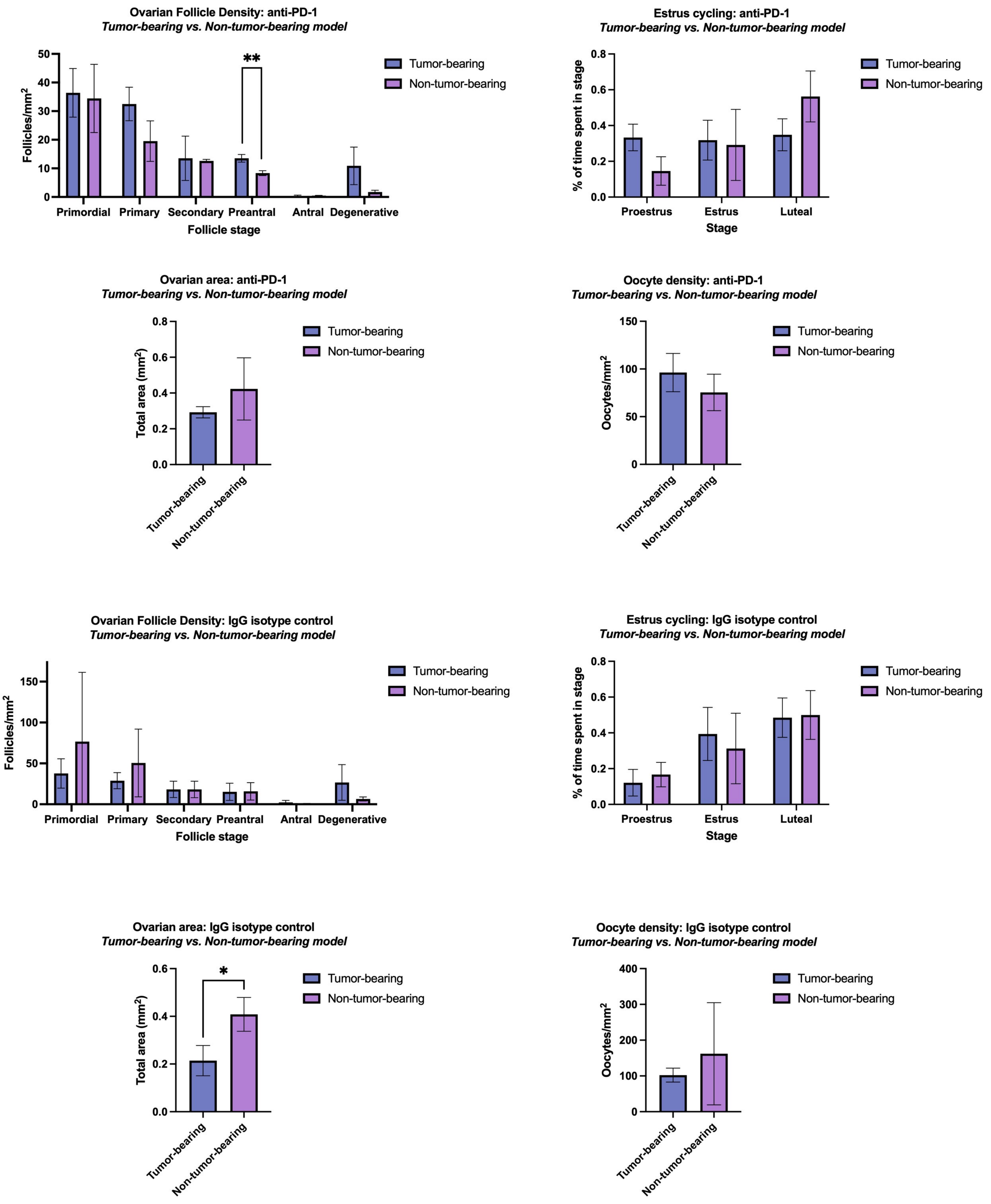
